# A Network of Phosphatidylinositol 4,5-bisphosphate Binding Sites Regulate Gating of the Ca^2+^-activated Cl^−^ Channel ANO1 (TMEM16A)

**DOI:** 10.1101/625897

**Authors:** Kuai Yu, Tao Jiang, YuanYuan Cui, Emad Tajkhorshid, H. Criss Hartzell

**Affiliations:** Department of Cell Biology, Emory University School of Medicine, Atlanta, GA, USA; NIH Center for Macromolecular Modeling and Bioinformatics, Beckman Institute for Advanced Science and Technology, Department of Biochemistry, and Center for Biophysics and Quantitative Biology, University of Illinois at Urbana-Champaign, Urbana, IL, USA

**Keywords:** chloride channel, protein-lipid interaction, phospholipid, structure-function, molecular dynamics

## Abstract

ANO1 (TMEM16A) is a Ca^2+^-activated Cl^−^ channel that regulates diverse cellular functions including fluid secretion, neuronal excitability, and smooth muscle contraction. ANO1 is activated by elevation of cytosolic Ca^2+^ and modulated by phosphatidylinositol 4,5-bisphosphate (PI(4,5)P_2_). Here we describe a closely concerted experimental and computational study, including electrophysiology, mutagenesis, functional assays, and extended sampling of lipid-protein interactions with molecular dynamics (MD) to characterize PI(4,5)P_2_ binding modes and sites on ANO1. ANO1 currents in excised inside-out patches activated by 270 nM Ca^2+^ at +100 mV are increased by exogenous PI(4,5)P_2_ with an EC_50_ = 1.24 µM. The effect of PI(4,5)P_2_ is dependent on membrane voltage and Ca^2+^ and is explained by a stabilization of the ANO1 Ca^2+^-bound open state. Unbiased atomistic MD simulations with 1.4 mol% PI(4,5)P_2_ in a phosphatidylcholine bilayer identified 8 binding sites with significant probability of binding PI(4,5)P_2_. Three of these sites captured 85% of all ANO1 - PI(4,5)P_2_ interactions. Mutagenesis of basic amino acids near the membrane-cytosol interface found three regions of ANO1 critical for PI(4,5)P_2_ regulation that correspond to the same three sites identified by MD. PI(4,5)P_2_ is stabilized by hydrogen bonding between amino acid sidechains and phosphate/hydroxyl groups on PI(4,5)P_2_. Binding of PI(4,5)P_2_ alters the position of the cytoplasmic extension of TM6, which plays a crucial role in ANO1 channel gating, and increases the accessibility of the inner vestibule to Cl^−^ions. We propose a model consisting of a network of three PI(4,5)P_2_ binding sites at the cytoplasmic face of the membrane allosterically regulating ANO1 channel gating.

**Significance statement:** Membrane proteins dwell in a sea of phospholipids that not only structurally stabilize the proteins by providing a hydrophobic environment for their transmembrane segments, but also dynamically regulate protein function. While many cation channels are known to be regulated by phosphatidylinositol 4,5-bisphosphate (PI(4,5)P_2_), relatively little is known about anion channel regulation by phosphoinositides. Using a combination of patch clamp electrophysiology and atomistic molecular dynamics simulations, we have identified several PI(4,5)P_2_ binding sites in ANO1 (TMEM16A), a Cl^−^ channel that performs myriad physiological functions from epithelial fluid secretion to regulation of electrical excitability. These binding sites form a band at the cytosolic interface of the membrane that we propose constitute a network to dynamically regulate this highly allosteric protein.

## Introduction

Ca^2+^-activated Cl^−^ channels (CaCCs) are jacks of all trades and masters of many. These ion channels facilitate the passive flow of Cl^−^ across cell membranes in response to elevation of cytosolic Ca^2+^ (1). Although CaCCs are probably best known for driving fluid secretion across mammalian epithelia, they are intimately involved in manifold physiological functions in all eukaryotes. CaCCs mediate action potentials in algae, the fast block to polyspermy in Anuran eggs, and regulate functions as diverse as smooth muscle contraction, nociception, neuronal excitability, insulin secretion, and cell proliferation and migration in mammals (1–13). While there are several types of CaCCs, the so-called classical CaCCs are encoded by the *ANO1* (*TMEM16A*) and *ANO2* (*TMEM16B*) genes (14–16).

Activation of ANO1 in its physiological context typically begins with ligand binding to a G-protein coupled receptor that activates phospholipase-C (PLC) (17–20). PLC hydrolyzes phosphatidylinositol 4,5-bisphosphate (PI(4,5)P_2_) in the plasma membrane to produce diacylgycerol and inositol 1,4,5-trisphosphate (Ins(1,4,5)P_3_). Ins(1,4,5)P_3_ binding to Ins(1,4,5)P_3_ receptors then triggers release of Ca^2+^ from internal ER stores and initiates store-operated Ca^2+^ entry. The resulting rise in cytosolic Ca^2+^ activates Cl^−^ flux because of conformational changes induced by direct binding of Ca^2+^ to the ANO1 protein (21–25). Calmodulin is not required for channel activation (21, 26).

Structurally, ANO1 is a dimer with each subunit composed of 10 transmembrane **(TM)** segments. The Cl^−^ selective pore of each subunit is surrounded by TMs 4–7 (22, 23, 27, 28) and each subunit has a Ca^2+^ binding site formed by amino acids E654, E702, E705, E734, and D738 in TMs 6– 8 (22-25, 29). Channel gating involves conformational changes of TM6 (24, 25, 30).

ANO1 is also regulated by PI(4,5)P_2_ (31–35). Ta et al. (33) have reported that PI(4,5)P_2_ stimulates ANO1 currents in excised patches. We (32) and Tembo et al. (35) have shown that PI(4,5)P_2_ can prevent and rescue Ca^2+^-dependent rundown caused by spontaneous PI(4,5)P_2_ hydrolysis. In whole-cell recording, we also showed that reduction of cellular PI(4,5)P_2_ by the voltage sensitive phosphatase Dr-VSP or by activation of GPCRs causes a reduction in ANO1 current (32). Pritchard et al. (34) show biochemical evidence that PI(4,5)P_2_ binds to ANO1, but they report that exogenous PI(4,5)P_2_ decreases endogenous Cl^−^currents thought to be encoded by *ANO1* in inside-out patches of pulmonary artery cells, in contrast to the results with heterologously-expressed ANO1 (32, 33).

PI(4,5)P_2_ binding sites typically have two or more positively charged amino acids with at least one Lys, and at least one aromatic residue (36–38). One well-known PI(4,5)P_2_ binding site is the pleckstrin homology (PH) domain which binds PI(4,5)P_2_ in a pocket of basic amino acids that are non-contiguous in the primary sequence but are close in proximity in the folded protein (39). On the other hand, “electrostatic type” binding sites typified by the CALM domain are clusters of contiguous basic amino acids that interact with phospholipids relatively non-specifically without forming a binding pocket (40). A wide variety of ion channels are regulated by PI(4,5)P_2_ (36-38, 41). In K_ir_ channels, these basic amino acids are located near the interface between the cytosol and the bulk lipid bilayer where the polar headgroups of lipids typically reside (42–44).

The objective of this study was to understand the mechanisms of regulation of ANO1 by PI(4,5)P_2_ and identify amino acids that stabilize PI(4,5)P_2_ binding. We find that PI(4,5)P_2_ stimulates ANO1 currents in excised patches in a voltage- and Ca^2+^-regulated manner. Mutagenesis and molecular dynamics (MD) simulations reveal that ANO1 has multiple PI(4,5)P_2_ binding sites. These data add to a growing body of knowledge showing that ANO1 is a highly allosteric protein that is gated by a network of interactions involving both Ca^2+^ and PI(4,5)P_2_.

## Results

### PI(4,5)P_2_ is a Positive Regulator of ANO1 Currents

To determine the effect of PI(4,5)P_2_ on ANO1, we transiently expressed ANO1 in HEK293 cells and measured the effect of dioctanoyl phosphatidylinositol 4,5-bisphosphate (diC8-PI(4,5)P_2_), a water-soluble, short acyl chain PI(4,5)P_2_, on inside-out excised patches. To reduce variability caused by different amounts of endogenous PI(4,5)P_2_, we first removed PI(4,5)P_2_ from the membrane patch by excising it into a solution containing 10 mM Mg^2+^ and 0 Ca^2+^/5 mM EGTA. Mg^2+^ competes with ANO1 for binding to PI(4,5)P_2_ by electrostatically masking negative charges on PI(4,5)P_2_ (45). ANO1 currents were measured during voltage steps first in the presence of 270 nM Ca^2+^ (I_initial_) and then with 270 nM Ca^2+^ plus 10 µM diC8-PI(4,5)P_2_ (I_PIP2_) (Fig. 1A). The increase in current (ΔI_PIP2_) was calculated as (I_PIP2_/I_initial_-1)*100%. On average, application of diC8-PI(4,5)P_2_ at +100 mV increased the current 79.1 ± 15.1% (n = 8). The effect of diC8-PI(4,5)P_2_ was slowly reversible: the currents decreased in amplitude when the patch was exposed to 270 nM Ca^2+^ in the absence of diC8-PI(4,5)P_2_ and exposure to 10 mM Mg^2+^ reduced the current further to its initial level (Fig. 1A).

**Figure 1.**
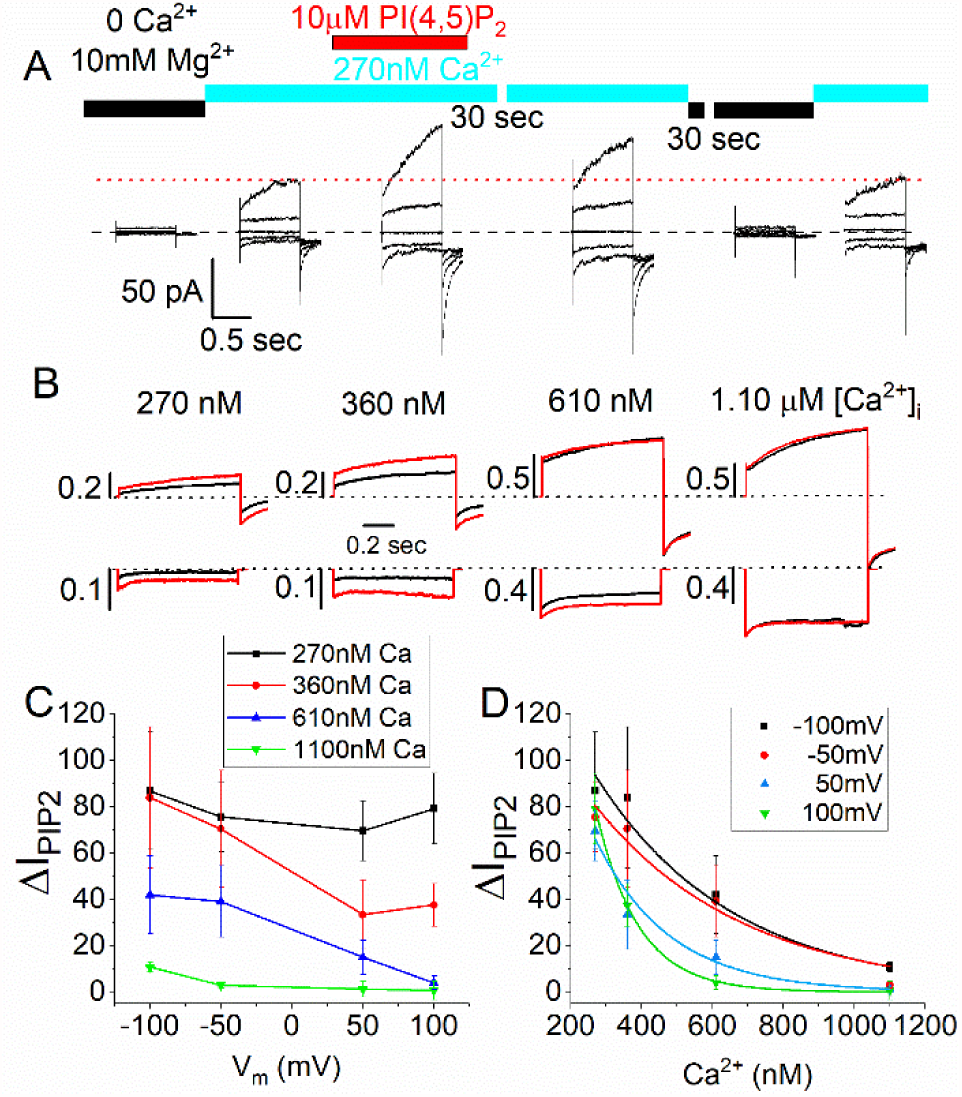
PI(4,5)P_2_ stimulates ANO1 current. **(A)** Current traces from a single inside-out excised patch exposed sequentially to solutions indicated above each set of traces. Each set of traces was obtained by voltage pulses from a holding potential of 0 mV to −100 mV, - 50 mV, 0 mV, 50 mV, and 100 mV. **(B)** Representative current traces from different patches exposed to different Ca^2+^ concentrations. Black trace: control solution without PI(4,5)P_2_. Red trace: same solution with added 10 µM diC8-PI(4,5)P_2_. Top row: +100 mV. Bottom row: −100 mV. Scale bars represent current amplitudes normalized to the current obtained from the same patch exposed to 20 µM Ca^2+^ to evoke a maximal current. **(C)** Voltage-dependence of PI(4,5)P_2_ stimulation of ANO1 current. Percent change in current amplitude caused by 10 µM diC8-PI(4,5)P_2_ (ΔI_PIP_2__) is plotted vs. voltage for patches exposed to 270 nM (black squares), 360 nM (red circles), 610 nM (blue triangles), or 1.1 µM (green inverted triangles) free Ca^2+^. **(D)** Effect of free Ca^2+^ on stimulation of ANO1 current by diC8-PI(4,5)P_2_. Percent change in current amplitude is plotted vs. free Ca^2+^ concentration at −100 mV (black squares), −50 mV (red circles), 50 mV (blue triangle), 100 mV (green inverted triangle).

### The Effect of PI(4,5)P_2_ is Regulated by Voltage and Ca^2+^

The effect of diC8-PI(4,5)P_2_ was modulated by both Ca^2+^ and voltage (Fig. 1B-D). At all voltages, ΔI_PIP2_ was greatest at lower Ca^2+^ concentrations (Fig. 1D). With 270 nM Ca^2+^, ΔI_PIP2_ at +100 mV was 79.1%, while with 1.1 µM Ca^2+^ ΔI_PIP2_ was <10%. Moreover, at all Ca^2+^ concentrations except the lowest [Ca^2+^] tested (270 nM), current stimulation by PI(4,5)P_2_ decreased with depolarization (Fig. 1C). For example, with 360 nM Ca^2+^ at −100 mV ΔI_PIP2_ was 83.8 ± 30%, but at +100 mV ΔI_PIP2_ was only 37.5 ± 9.1%. The IC_50_ for Ca^2+^ suppression of the PI(4,5)P_2_ effect was determined by fitting the data in Fig. 1D to exponential equations. The estimated IC_50_ for attenuation of the diC8-PI(4,5)P_2_ stimulatory effect was 390 nM Ca^2+^ at 100 mV and 656 nM Ca^2+^ at −100 mV (Fig. 1D). Because ANO1 has a lower affinity for Ca^2+^ at hyperpolarized potentials (46), the observation that PI(4,5)P_2_ has a larger effect at hyperpolarized potentials and at lower Ca^2+^ concentrations indicates that the lipid stabilizes the open, Ca^2+^-liganded state of the channel. At saturating Ca^2+^ concentration, PI(4,5)P_2_ has no additional effect.

### The Effect of PI(4,5)P_2_ is Specific and Moderate Affinity

We compared the effects of two other physiologically important phosphoinositides (dioctanoyl phosphatidylinositol 3,4,5-trisphosphate [diC8-PI(3,4,5)P_3_] and dioctanoyl phosphatidylinositol 3,5-bisphosphate [diC8-PI(3,5)P_2_]) (47, 48) on ANO1 currents (Fig. 2). diC8-PI(3,4,5)P_3_, which has one additional phosphate at the 3-position, was less than half as efficacious at 10 µM as diC8-PI(4,5)P_2_: ANO1 current increased 31 ± 12 % (Fig. 2B,E,G). diC8-PI(3,5)P_2_, which has a phosphate in the 3-position instead of the 4-position had no effect at 10 µM (Fig. 2C,F,G). Thus, the effect of PI(4,5)P_2_ on ANO1 exhibits specificity.

**Figure 2.**
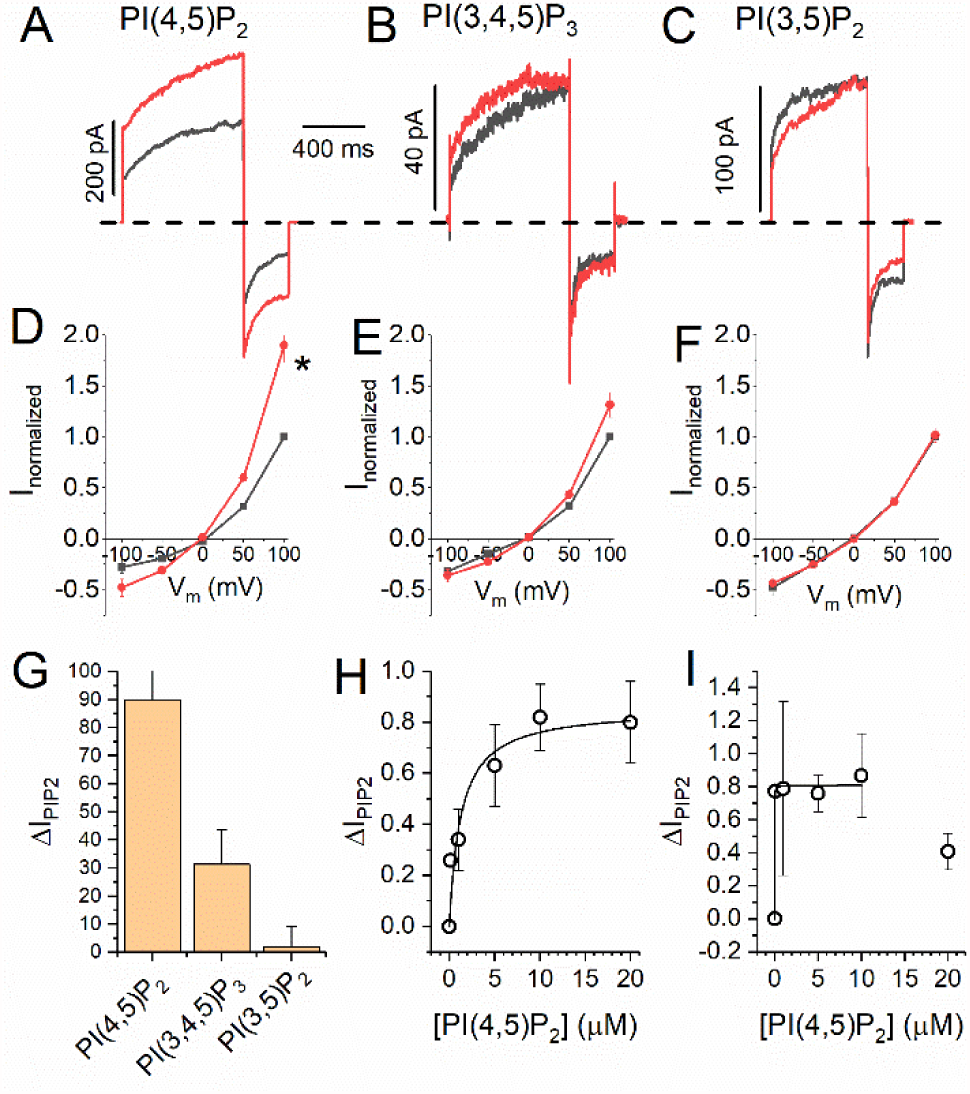
PI(4,5)P_2_ effect is selective and moderately high affinity. **(A-C)** Representative current traces at +100 mV with 270 nM Ca^2+^ in the absence (black) and presence (red) of 10 µM **(A)** diC8-PI(4,5)P_2_, **(B)** diC8-PI(3,4,5,6)P_3_, **(C)** PI(3,5)P_2_. **(D-F)** Average ANO1 current voltage relationships in the absence of phosphoinositides (black squares) and in the presence of 10 µM **(A)** diC8-PI(4,5)P_2_, **(B)** diC8-PI(3,4,5,6)P_3_, **(C)** PI(3,5)P_2_. **(G)** Summary of stimulatory effect of phosphoinositides on ANO1 current at 100 mV with 270 nM Ca^2+^. **(H-I)** Concentration-response curve for diC8-PI(4,5)P_2_ on ANO1 current with 270 nM Ca^2+^at **(H)** 100 mV and **(I)** −100 mV.

To estimate the apparent EC_50_ for PI(4,5)P_2_, we measured ΔI_PIP2_ in response to different diC8-PI(4,5)P_2_ concentrations. The data, fitted to the Michaelis-Menton equation, gave an EC_50_ = 1.24 µM (Fig. 2H) at 100 mV with 270 nM Ca^2+^. This falls within the range for the effect of PI(4,5)P_2_ on other ion channels (0.12 - 4.6 µM, (49–51)). At −100 mV, inward currents were more sensitive to diC8-PI(4,5)P_2_ with EC_50_ < 0.1 µM (Fig. 2I).

### Selection of Potential PI(4,5)P_2_ Regulatory Sites

To identify amino acids that might play a role in PI(4,5)P_2_ regulation of ANO1, we mutagenized amino acids singly or in groups and tested whether stimulation of ANO1 currents in excised patches by diC8-PI(4,5)P_2_ was altered. We focused on basic amino acids because known PI(4,5)P_2_ binding sites contain two or more positively charged amino acids (38, 52). There are 62 Lys and 60 Arg residues in mouse ANO1. To narrow candidate residues for mutagenesis, we identified Lys and Arg residues located within 10 Å of the cytoplasmic membrane interface. To this list, we added amino acids in the cytoplasmic N-terminus: K71, R72, R81, and R82 because they are predicted to constitute a phospholipid binding site by BHSEARCH (53), and K124, R125, R127, and R128 because they align partly with a region that ostensibly forms a PI(4,5)P_2_ binding site (54, 55) in the ANO1 paralog ANO6 (amino acids 93 – 100) (54, 55) (Fig. 3A).

**Figure 3.**
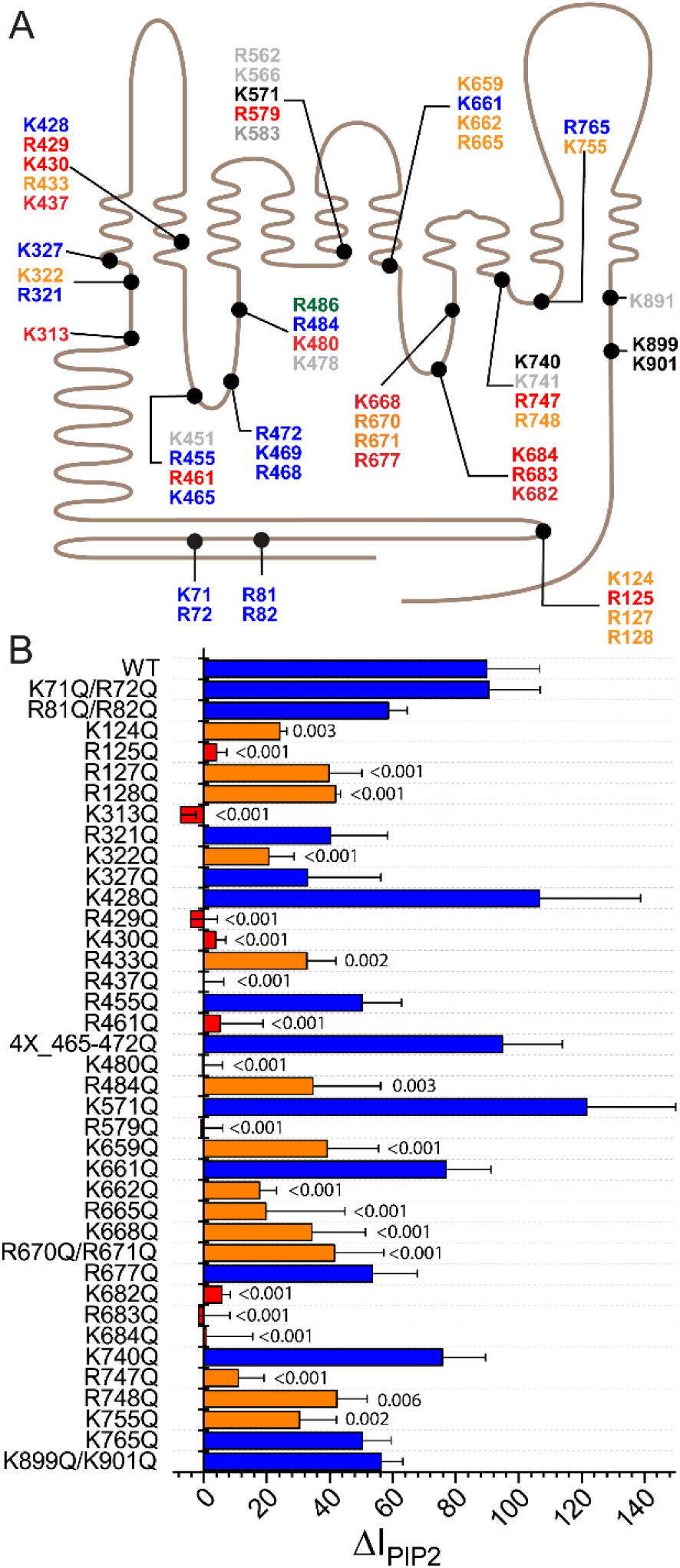
Amino acids involved in mediating the effect PI(4,5)P_2_ on ANO1. **(A)** Schematic of 51 basic amino acids that were considered potential PI(4,5)P_2_ interacting residues. ANO1 is represented as a line with 10 transmembrane helices. Extracellular is upwards. The length of the line is scaled to the amino acid sequence. Colors indicate effect of mutagenesis on stimulation of ANO1 current by PI(4,5)P_2_ corresponding to panel B: Black, response was like WT; Red, PI(4,5)P_2_ effect was essentially abolished; Orange: PI(4,5)P_2_ effect was significantly reduced; Grey: not tested. **(B)** Effects of mutation of basic amino acids in ANO1 on the stimulation of ANO1 current in inside-out patches by PI(4,5)P_2_. 10 µM PI(4,5)P_2_ stimulates ANO1 current by ΔI_PIP2_ = 89.9%. Error bars are ± S.E.M. n = 3 −13 patches per mutation. Numbers above bars show statistical P calculated by one-way ANOVA with Fisher LSD post-hoc analysis for difference between means. Blue bars: P >0.05. Orange bars: P <0.05 and >50% reduction in response to PI(4,5)P_2_. Red bars: P <0.01 and >90% reduction in response to PI(4,5)P_2_.

### Mutagenesis to Identify PI(4,5)P_2_ Regulatory Sites

We neutralized the charge on selected Arg and Lys residues by replacement with Gln. Surprisingly, more than 20 mutants were found to have a statistically significant reduction in response to diC8-PI(4,5)P_2_ compared to WT. For WT ANO1, ΔI_PIP2_ = 89.9 ± 16.8% (n = 13) in this set of experiments. ΔI_PIP2_ for “significant” mutants was <40% (p < 0.05, Fig. 3B orange and red bars). Eleven amino acids were considered “critical” for PI(4,5)P_2_ regulation of ANO1 because their mutation reduced ΔI_PIP2_ to <10% (Fig. 3B, red bars).

These critical residues define three locations in the ANO1 structure that correspond to pockets of surface electronegativity (Fig. 4). In the following description, the number in parenthesis is ΔI_PIP2_ when this amino acid is substituted with Gln (WT = 89.9%). Site A is near the dimer interface and is defined by critical amino acids R429 (−4%), K430 (4%), and R437 (0%) in TM2 and K313 (−8%) preceding TM1. R433 (33%), located one helix turn from K430, also contributes to this site. Site B is located at the cytoplasmic end of TM6, which plays a central role in ANO1 gating. It is defined by K682 (6%), R683 (−2%), and K684 (1%). Three nearby amino acids also significantly reduce the effect of PI(4,5)P_2_: K662 (18%), K668 (38%), and K665 (22%). Site C is located in the short intracellular loop between TM2 and TM3 that forms one side of the Cl^−^ ion permeation pathway. It is defined by R461 (6%), K480 (0%), and R484 (39%).

**Figure 4.**
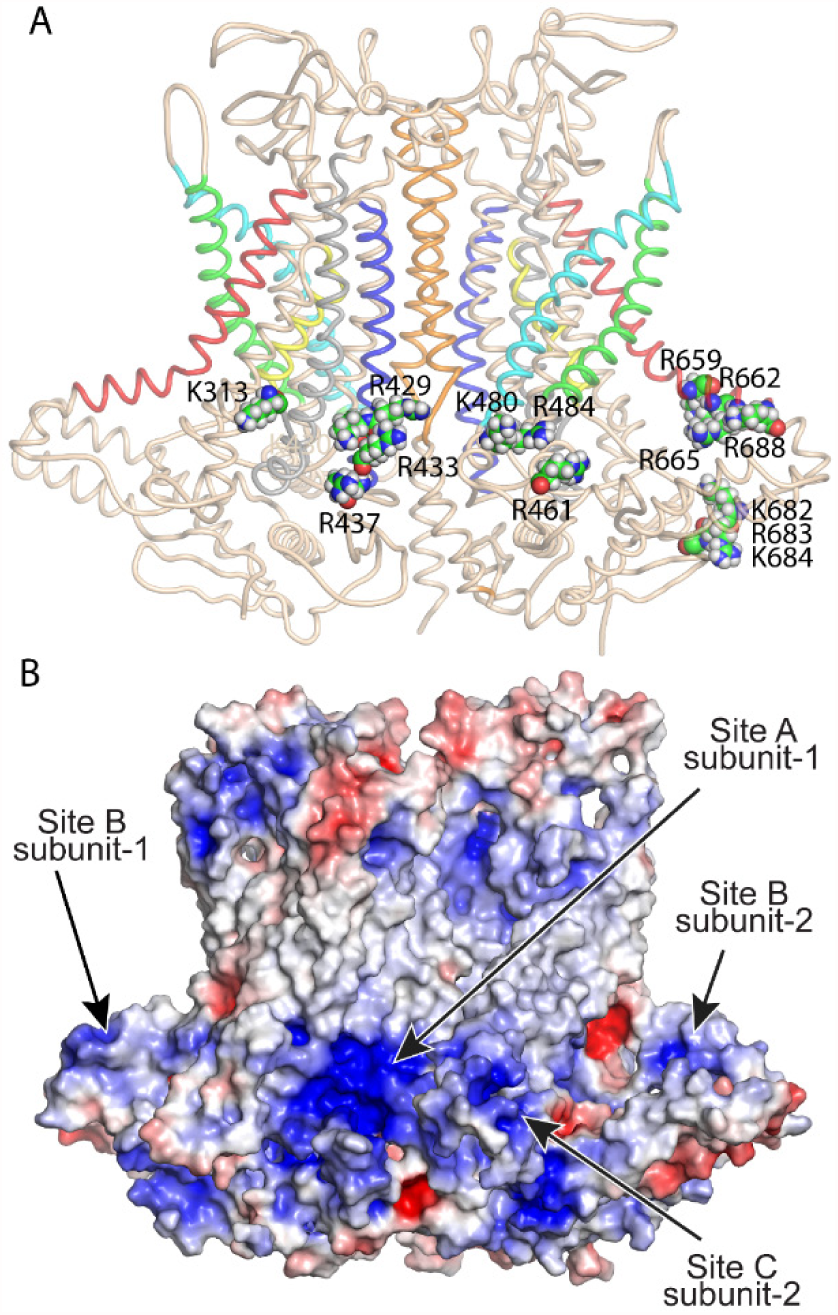
Location of mutations that are critical for the stimulatory effect of PI(4,5)P_2_ on ANO1. **(A)** Cartoon representation of ANO1 with critical amino acids as space-fill labeled. For clarity, Site A (K313, R429, K430, R433, R437) is shown only in the left subunit; Site B (K659, R662, R665, R668, R682, R683, and K684) and Site C (R461, K480, and R484) are shown only in the right subunit. Helices are colored, blue (TM2), cyan (TM3), green (TM4), red (TM6), yellow (TM7), and orange (TM10). **(B)** Electrostatic surface of ANO1 calculated by APBS Electrostatics in PYMOL. Red (+5 kT/e), Blue (−5 kT/e).

### Microscopic Characterization of PI(4,5)P_2_ – ANO1 Interaction

In order to gain more insight into the binding of PI(4,5)P_2_ t o ANO1, we performed molecular dynamics (MD) simulations using the highly mobile membrane mimetic model (HMMM) (54,59). The HMMM model was introduced to accelerate lipid diffusion in atomistic MD simulations in order to obtain significantly enhanced sampling of interaction of lipid headgroups with proteins within simulation timescales currently achievable. In conventional membrane models, lipids with long acyl chains have a relatively low lateral diffusion constant (∼8 × 10^−8^ cm^2^ s^−1^). Therefore, during a 500 ns simulation, lipids do not diffuse far enough to allow sufficient mixing and sampling of the protein surface. The HMMM model accelerates lipid lateral diffusion by shortening lipid acyl tails to six carbons and filling the membrane core with a liquid phase (Fig. 5A). The HMMM model has been used successfully to study the diffusion and domain formation of lipids and membrane-associated proteins, pore formation by peptides, hydrophobic matching of transmembrane helices, and membrane-associated assembly of coagulation factors (56–64). The purpose of the model is to provide a more flexible and mobile environment that allows for rapid rearrangement and displacement of the lipid headgroups, thereby facilitating phenomena that might be inaccessible with conventional membrane models due to the inherently slow dynamics of the lipids.

**Figure 5.**
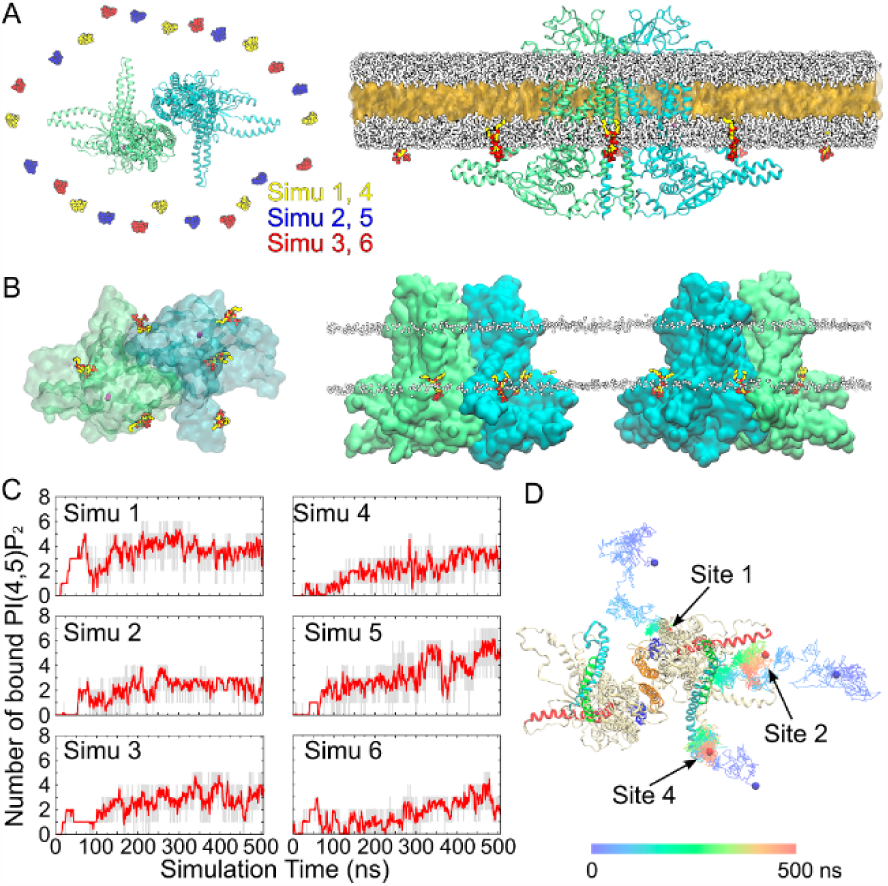
Spontaneous PI(4,5)P_2_ binding captured in MD simulations. **(A)** (left) Top view showing the initial placement of eight PI(4,5)P_2_ molecules in the inner leaflet of the bilayer surrounding the channel. Six independent simulations were performed with PI(4,5)P_2_ molecules placed in different initial positions, as shown by different colors. The two subunits of ANO1 are shown in green and blue, respectively. (right) The initial HMMM simulation system viewed from the membrane, in which a large fraction of the acyl tails of the membrane-forming lipids is replaced by a liquid organic phase (yellow surface representation). The short-tailed POPC molecules are shown in white. The short-tailed PI(4,5)P_2_ molecules are shown in color (oxygen: red, phosphorus: tan, carbon: yellow). Ions and water molecules are not shown for clarity. **(B)** Top and side views showing the positions of the PI(4,5)P_2_ molecules at the end of one representative HMMM simulation (simu 5). Six PI(4,5)P_2_ molecules were found to be directly coordinated by the charged/polar residues of the channel. Ca^2+^ ions are shown as purple spheres. The membrane position is demarcated by the phosphorus atoms (white spheres) of the PC lipids. **(C)** Number of bound PI(4,5)P_2_ molecules (grey trace) and the moving average (red, bin=20) plotted vs. simulation time for the six simulations. **(D)** Representative trajectories showing the binding of PI(4,5)P_2_ to Sites 1, 2, and 4. The colored lines illustrate the position of the 4’-phosphate of PI(4,5)P_2_ over the 500-ns simulations at 100-ps intervals. The initial and final positions of the 4’-phosphate are shown as blue and red spheres, respectively.

To examine the interaction of ANO1 with PI(4,5)P_2_, eight C6-PI(4,5)P_2_ molecules were added to the inner leaflet of a C6-POPC bilayer evenly surrounding ANO1 (Fig. 5A) at the beginning of the simulation. C6-PI(4,5)P_2_ constitutes ∼1.4 mol% of the total lipids in the inner leaflet. To diminish any bias introduced by the initial placement, the position of the PI(4,5)P_2_ molecules were shifted in each of the six independent simulations such that each simulation began with different PI(4,5)P_2_ starting positions. Within 50 ns, the PI(4,5)P_2_ molecules diffused around the protein and interacted with it. The number of bound PI(4,5)P_2_ molecules reached a plateau after ∼300 ns with an average of 4.5 PI(4,5)P_2_ molecules bound per dimer (range 3 – 6) in each simulation at the end of 500 ns (Fig. 5 B,C). Representative trajectories of PI(4,5)P_2_ interacting with ANO1 are shown in Fig. 5D. At the end of the six HMMM simulations with C6-PI(4,5)P_2_, all subunits had at least one PI(4,5)P_2_ molecule bound: two subunits had Site 1 occupied only, six subunits had Site 2 and/or 4 occupied, 4 subunits had Site 1 plus Site 2 and/or 4 occupied.

By calculating the density of the inositol groups over the simulation trajectories and the interactions between the phosphate/hydroxyl groups of PI(4,5)P_2_ and ANO1, eight sites on the cytoplasmic surface of ANO1 were found to be visited by PI(4,5)P_2_ molecules. Sites 1–5 (labelled according to their clockwise location) involve basic and aromatic residues at the membrane-cytoplasm interface around the ends of transmembrane helices TM1, TM2, TM3, TM4, TM6, and TM10 (Fig. 6A). PI(4,5)P_2_ binding to these sites was observed in multiple independent trajectories and for both subunits of ANO1 (Fig. 6B), revealing several specific and stable interactions between PI(4,5)P_2_ and the protein. Sites 6–8, which bind PI(4,5)P_2_ only transiently, mainly involve residues from the N-terminal region of ANO1, including the short α-helices α0a and α0b and the loop preceding them, and the loop between Nβ1 and Nα1. Among the eight sites captured in the simulations, Sites 1, 2, and 4 showed the highest occupancies (Fig. 6C). These three sites represent 84.3% of all the binding events observed. PI(4,5)P_2_ headgroups are well coordinated in these binding sites, resulting in residence lifetimes as long as 116.5 ns (Fig. 6D). Because PI(4,5)P_2_ binding to Sites 1, 2, and 4 was more robust than binding to the other sites, we focused on these three sites for further analysis.

**Figure 6.**
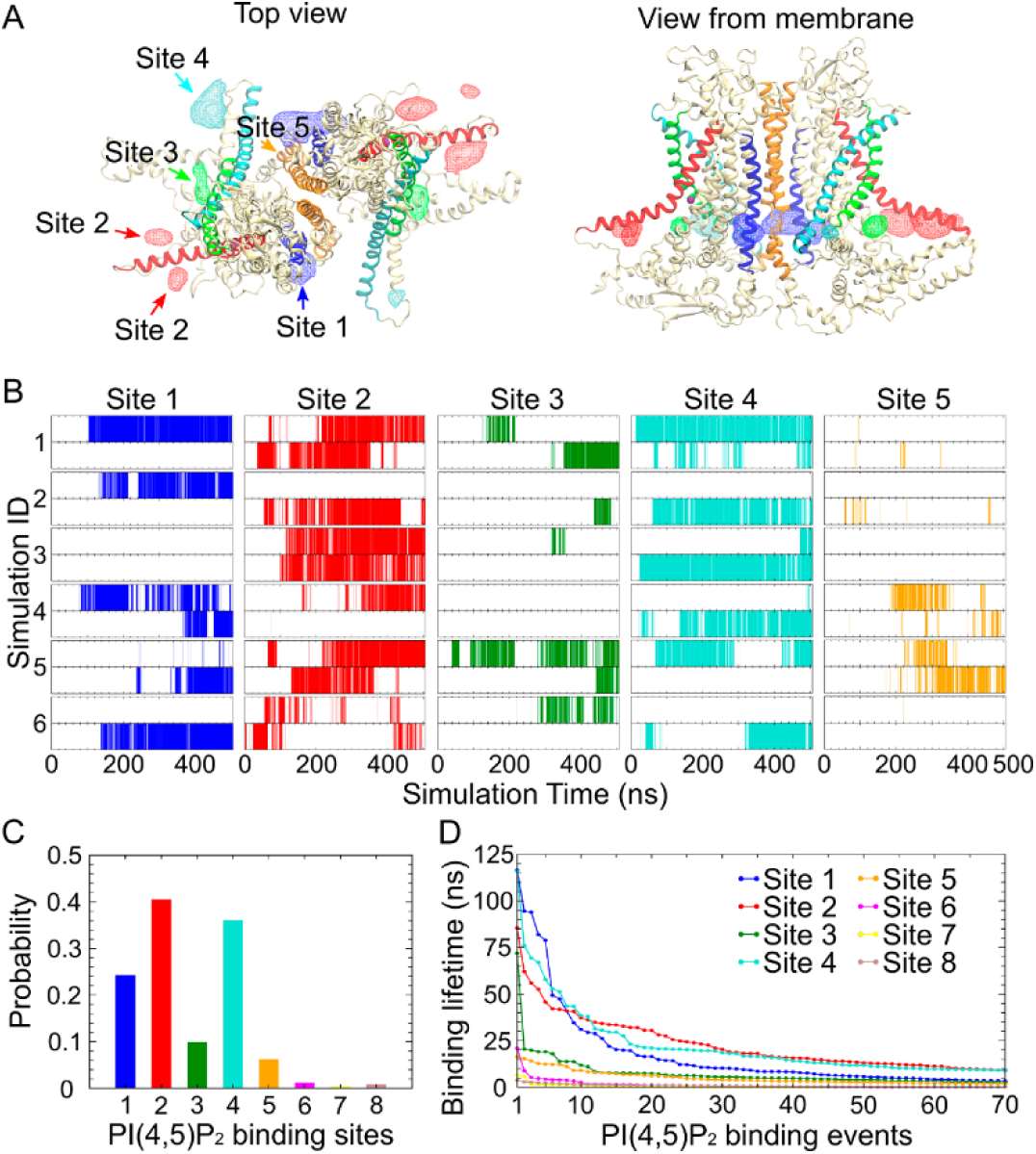
PI(4,5)P_2_ binding at specific sites on ANO1. **(A)** Volumetric map of inositol occupancy extracted from the last 200 ns of the HMMM simulations is shown as colored wireframe contoured at isovalue 0.05 overlaid on the protein structure. Analysis combines results from the six simulations. Each local map is colored with the same color used for the nearby transmembrane helix that is involved in PI(4,5)P_2_ binding (TM2: blue, TM3: cyan, TM4: green, TM6: red, and TM10: orange). **(B)** PI(4,5)P_2_ occupancy at Sites 1–5 in each subunit (upper panel: Subunit 1, lower panel: Subunit 2) during the 6 independent PI(4,5)P_2_-binding simulations. PI(4,5)P_2_ binding is indicated by the vertical lines at the corresponding time point. **(C)** The probability of PI(4,5)P_2_ occupancy for each site over the 6 PI(4,5)P_2_-binding simulations. PI(4,5)P_2_ binding was observed mainly in Sites 1, 2, and 4. **(D)** Dwell time of the top 70 binding events in each of the 8 binding sites, sorted in descending order. Only Sites 1, 2, and 4 exhibit significant dwell times.

### Insight Into the Functional Roles of Specific Key Amino Acids

In Site 1, five basic residues (R433, K430, R429, R437, and K313, Fig. 7A,D), which stabilize the binding of PI(4,5)P_2_ by coordinating the inositol ring and its phosphates. All of these residues are experimentally verified to strongly affect PI(4,5)P_2_ binding (Fig. 3B). Of the four residues that coordinate PI(4,5)P_2_ for >10% of the total residence time in Site 1, K430 shows strong preference for interaction with the hydroxyl groups on the inositol ring, while the other three basic residues mainly coordinate 4’- and 5’-phosphate groups. R437 is located closer to the cytoplasm than the other residues and interacts almost exclusively with the 4’- and 5’-phosphate groups. This feature suggests that R437 might be crucial for specific recognition of PI(4,5)P_2_. Experimentally, the R437Q mutation totally abolishes the PI(4,5)P_2_-induced effect. Similarly, in Site 2 (Fig. 7B,E), all the residues facing the cytoplasm including those in the cytoplasmic loop connecting TM6 and TM7 (R677, R683, K682, K684) and R665 in TM6 are mainly involved in coordinating the 4’- and 5’-phosphate groups. The other residues in TM6 show less discrimination and also interact with the 1’-phosphate and hydroxyl groups on the inositol ring. In Site 4 (Fig. 7C,F), all major interacting residues show preference for 4’- and 5’-phosphate groups except for R484, which indiscriminately coordinates all the functional groups on the inositol ring.

**Figure 7.**
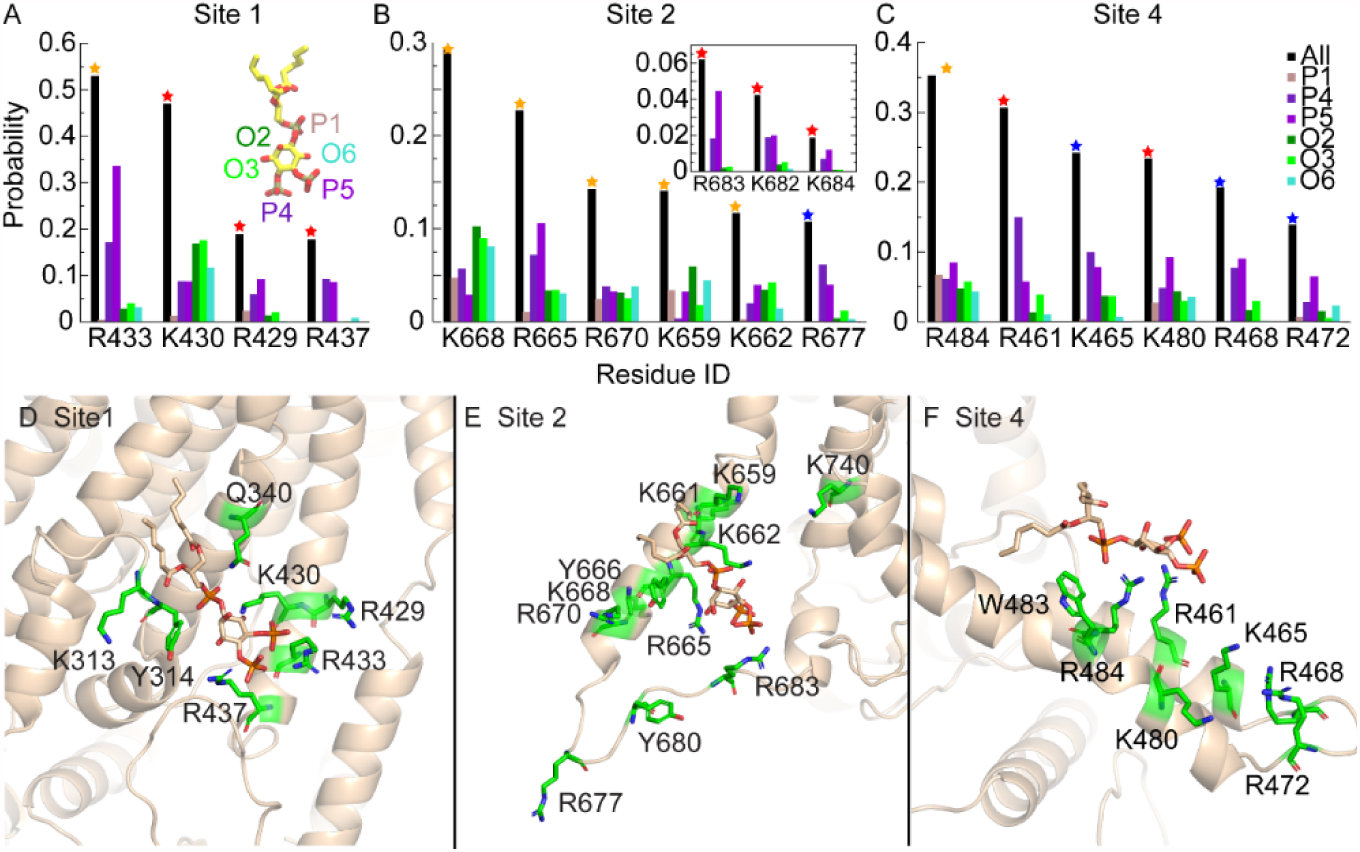
PI(4,5)P_2_ coordination in major binding sites. **(A-C)** The probability of PI(4,5)P_2_ headgroup coordination by the key basic residues in Sites 1, 2, and 4 is shown as black bars. Residues that coordinate PI(4,5)P_2_ for >10% of the total binding time in each site are shown (other residues that affect PI(4,5)P_2_ binding experimentally are also shown in the inset). The color of the star on each bar represents the experimental results in Figure 3(B), where the mutation of the residues affects PI(4,5)P_2_ binding to different degrees. The coordination probability for each functional group on the inositol ring (1’-phosphate, 4’-phosphate, 5’-phosphate, 2’-hydroxyl, 3’-hydroxyl, and 6’-hydroxyl) is shown individually. **(D-F)** Coordination of PI(4,5)P_2_ in Sites 1, 2, and 4. Amino acids that interact with each binding site are shown as stick representation in green. PI(4,5)P_2_ is shown in stick representation in tan.

### Specificity of Phospholipid Binding

To evaluate the specificity of PI(4,5)P_2_ binding, we measured the effect of PtdSer on PI(4,5)P_2_ binding by performing three 350-ns simulations with a mixture 8 PI(4,5)P_2_ and 8 PtdSer molecules with each simulation having a different initial placement of the lipids. On average, the number of bound PI(4,5)P_2_ molecules over the trajectories was reduced ∼20% from 2.73 ± 1.28 to 2.21 ± 1.45 (per monomer), which was not statistically significant. In these simulations, PI(4,5)P_2_ interacted with 42 residues. We analyzed whether the PI(4,5)P_2_ coordination probability was different for these 42 residues in simulations with and without PtdSer and found that they were not statistically different (one way ANOVA, p>0.1). In contrast, PtdSer binding to these sites was significantly different than PI(4,5)P_2_ in the same simulations (p <0.001). In these simulations, the PtdSer headgroup only interacted with seven residues with significant probability (>0.01). Two residues are in Site 1 (R429: 0.09; R433: 0.06), one in Site 2 (R670: 0.02), two in Site 4 (R461: 0.02, K465: 0.01), and two others near Site 4 (R486: 0.04, K469: 0.01). These data suggest that although PtdSer may also bind to these sites, PI(4,5)P_2_ has a higher probability of interaction.

### Potential Conformational Changes Induced by PI(4,5)P_2_

To determine whether binding of PI(4,5)P_2_ was influenced by full-length acyl chains, the short-tailed lipid molecules (C6-PI(4,5)P_2_ and C6-POPC) used in the initial probing of lipid-protein interactions were converted back to full-length lipids (palmitoyl-oleoyl-PI(4,5)P_2_ and palmitoyl-oleoyl-POPC) at the end of the HMMM simulations, and the resulting full systems were subjected to an additional equilibrium simulation of 600 ns. During these simulations with full-length lipid, each bound PI(4,5)P_2_ molecule remained coordinated around its binding site, suggesting that these sites stably bind full-length PI(4,5)P_2_. In the HMMM simulations, the PI(4,5)P_2_ molecules were more likely to dissociate from their binding sites and sample more binding events.

We compared the conformation of the PI(4,5)P_2_-bound ANO1 with its conformation in a control simulation performed in the absence of PI(4,5)P_2_. The most significant change captured in the simulations with PI(4,5)P_2_ bound is a rotation of the cytoplasmic half of TM6, which forms one side of the channel pore and plays a key role in channel gating (Fig. 8A,B). The position of the cytoplasmic portion of TM6 is described by two angles. α is the angle of TM6 projected onto the x-y plane with the position in the cryo-EM structure defined as 0°. β is the angle of TM6 relative to the x-y plane. In the absence of PI(4,5)P_2_, the cytoplasmic end of TM6 fluctuates only slightly around its initial position (peak α = 0°, peak β = 36°). When any site is occupied at Sites 2/4, or single occupancy at Site 1, there is a significant shift of TM6 away from the pore (peak α = 12° – 18°, peak β = 22°). When Site 1 plus Site 2 and/or 4 are occupied, there is an even more dramatic rotation of the cytoplasmic end of TM6 away from the pore (peak α = 18° – 32°, peak β = 22°).

**Figure 8.**
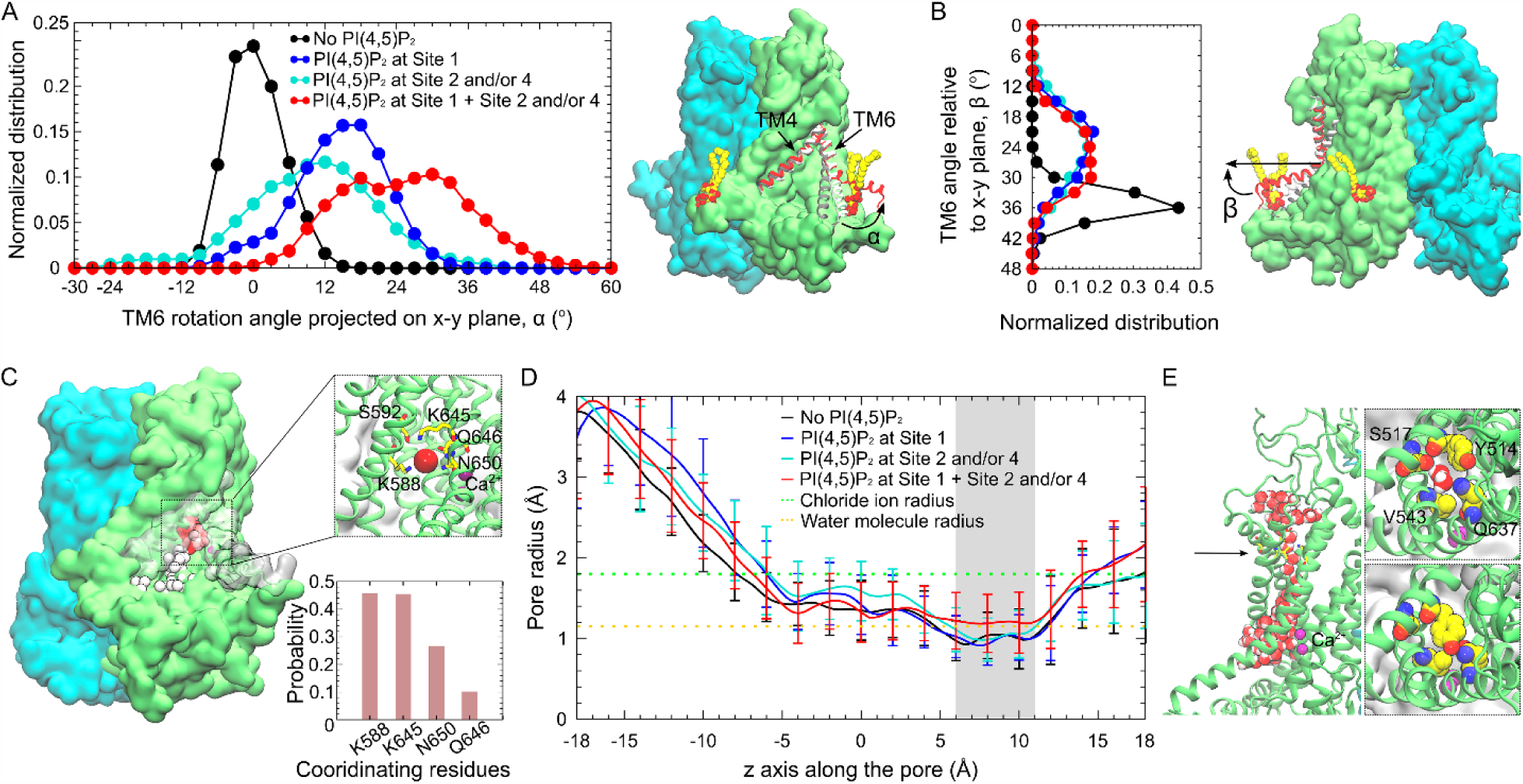
Conformational changes induced by PI(4,5)P_2_ binding. **(A)** Rotation angle of the cytoplasmic portion of TM6 with respect to its position in the cryo-EM structure, projected onto the x-y plane. Left panel: Distribution of the rotation angle normalized over the 600 ns full-length lipid simulations for each palmitoyl-oleoyl PI(4,5)P_2_ occupancy state. The cytoplasmic portion of TM6 fluctuates around its initial position (0°) in the absence of PI(4,5)P_2_ (black). Single occupancy of Site 1 (blue), or single/double occupancy of Sites 2 and 4 (cyan) shifts the rotation angle to positive values (cytoplasmic portion of TM6 away from the pore). Multiple occupancy of PI(4,5)P_2_ at Sites 1 plus 2 and/or 4 (red) rotates TM6 most dramatically away from the pore. Right panel: Snapshots of TM6 in the multiply-occupied state (red), the PI(4,5)P_2_-free state (black), and the cryo-EM structure (white). Bound PI(4,5)P_2_ is shown in vdW representation. **(B)** Left panel: Distribution of the angle between the TM6 cytoplasmic portion and the x-y plane. Right panel: Snapshot showing the angle relative to the membrane in the multiple-occupied state (red), the PI(4,5)P_2_-free state (black), and the cryo-EM structure (white). **(C)** Time series snapshots showing the spontaneous penetration of Cl^−^ ions (vdW spheres, white-to-red over time) in the multiply-occupied state. Coordination of the Cl^−^ ion deeply binding inside the pore is shown in the inset, with the probability of coordination for all the binding events shown below. Cl^−^ and Ca^2+^ ions are shown as red and purple spheres, respectively. **(D)** Average radius of the ion conduction pore calculated using HOLE (86) illustrates a bottleneck between 6 and 11 Å (grey shading). The bottleneck in the multiply-occupied state dilates ∼30% compared to other states. Orange dotted line: radius of water molecule (1.15 Å). Green dotted line: radius of Cl^−^ ion (1.8 Å). **(E)** Left panel: Hydration of the ion conduction pore when the bottleneck (black arrow) dilates in the multiply-occupied state. Right panel: Top views of the pore showing the amino acids forming the bottleneck. The sidechain orientation of Y514 affects the pore radius and hydration.

When PI(4,5)P_2_ was bound, penetration of Cl^−^ions into the pore was frequently observed (Fig. 8C). Especially in the case of the multi-occupied simulations (sites 1 and 2/4), deep penetration of Cl^−^ occurred ∼25% of the total time. The major Cl^−^-coordinating amino acids were K588, K645, N650, and Q646. S592 also participated when Cl^−^ entered more deeply. Although we had hoped to observe Cl^−^ ions transiting the entire conduction pathway, this did not happen in the timeframe of our simulations. However, these simulations were performed with no applied voltage. Without driving force for Cl^−^ movement, ion conduction may be too slow to capture in this 600 ns simulation. Further, the voltage-dependent gating of ANO1 was not activated, so the conditions for the protein entering a conducting state may be sub-optimal. The cryo-EM model of ANO1 we used for these simulations is a non-conducting state with a constriction in the pore between 6 and 11 Å formed by the bulky residue Y514 and three nearby residues (S517, V543, and Q637). When multiple PI(4,5)P_2_ sites are occupied (site 1 and 2/4), we observed that the conduction pathway dilated ∼30% to a radius of 1.2 Å (Fig. 8D,E), which is slightly greater than the radius of a water molecule (1.15 Å), but is smaller than a Cl^−^ ion (1.8 Å). Thus, forces that stabilize this non-conducting conformation are apparently sufficiently strong to prevent pore opening in the timeframe of these simulations.

## Discussion

### Correspondence between Mutagenesis and MD Predictions

In this study, we used mutagenesis and computational modeling to identify lipid-binding sites in ANO1 that are important in regulation of the channel by PI(4,5)P_2_. Sites 1, 2, and 4 predicted by MD simulations to capture 84% of all interactions between ANO1 and PI(4,5)P_2_ correspond topographically to Sites A, B, and C determined by mutagenesis. In further commentary, these sites are referred to as Sites A/1, B/2, and C/4.

Both the mutagenesis and computational approaches have their limitations. Mutagenesis experiments can identify sites that are important in functionally mediating the effect of PI(4,5)P_2_ on ANO1, but this approach cannot easily distinguish between amino acids that coordinate PI(4,5)P_2_ binding, stabilize the structure of the binding site, or allosterically couple PI(4,5)P_2_ binding to channel gating or ion permeation. Furthermore, it should be noted that amino acids critical for the PI(4,5)P_2_ effect were identified under very specific recording conditions of 270 nM Ca^2+^ and 100 mV. Given the sensitivity of the PI(4,5)P_2_ effect to both voltage and Ca^2+^, it is entirely possible that under different conditions where ANO1 occupies different conformational states, a different complement of amino acids may be involved in mediating PI(4,5)P_2_ binding and effect. The MD simulations provide direct information about amino acids that are energetically favorable for binding PI(4,5)P_2_, but suffer from incomplete knowledge about native ANO1 and membrane structure in living cells. Furthermore, the MD simulations were not performed under identical conditions as the live-cell patch clamp experiments. For example, no voltage was applied during the MD simulations and there was no free Ca^2+^ present in the system. While the correspondence between the functional live-cell experiments and the MD simulations is not perfect, the global agreement is remarkable. This provides a high level of confidence that these sites are involved in PI(4,5)P_2_ binding.

Site A/1 is the most robust site we have identified with all the same amino acids identified in mutagenesis and MD (Table I). Mutation of a single amino acid (R429, K430, R437, or K313) in this site nearly abolishes the effect of PI(4,5)P_2_. The MD simulation shows the headgroup of PI(4,5)P_2_ is well-coordinated by a pocket formed by R429, K430, and R437 on one side and K313, Y314, and Q340 on the other sides (Fig. 7D). The site fulfils the requirements for a canonical PI(4,5)P_2_ binding site by having at least one Lys (K430) and one aromatic amino acid (Y314).

**Table I.** Comparison of amino acids in PI(4,5)P_2_ sites identified by MD simulation and mutagenesis. MD probability is probability of the amino acid coordinating PI(4,5)P_2_ within that site (Fig. 7). PI(4,5)P_2_ effect is expressed as % of effect on wild type ANO1.

|  | MD Simulation | PI(4,5)P<sub>2</sub> Effect |
| --- | --- | --- |
|  | Probability | Mutant/WT X100 |
| **SITE A/1** |  |  |
| R433 | 0.52 | 36 |
| K430 | 0.47 | 4 |
| R429 | 0.18 | -5 |
| R437 | 0.17 | 0 |
| K313 | 0.07 | -8 |
| **SITE B/2** |  |  |
| K668 | 0.29 | 42 |
| R665 | 0.22 | 23 |
| R670 | 0.15 | 46 |
| K659 | 0.15 | 44 |
| K662 | 0.13 | 20 |
| R677 | 0.12 | 60 |
| K740 | 0.06 | 84 |
| R683 | 0.06 | -2 |
| K661 | 0.05 | 86 |
| K682 | - | 6 |
| K684 | - | 1 |
| **SITE C/4** |  |  |
| R484 | 0.35 | 48 |
| R461 | 0.31 | 10 |
| K465 | 0.25 | 106 |
| K480 | 0.23 | 0 |
| R468 | 0.20 | 100 |
| R472 | 0.14 | 100 |
| **OTHERS** |  |  |
| R125 | N-terminus | 4 |
| R579 | Structural ? | -1 |
| R747 | Structural ? | 12 |

A close agreement also exists between the MD and functional studies for Site B/2 (Table I). By both MD and mutagenesis, Site B/2 is composed of at least six basic amino acids, only one of which (R683) is essential for the PI(4,5)P_2_ effect. Mutation of each of the other amino acids has a partial effect, but this agrees with the modest probability of each of these amino acids coordinating PI(4,5)P_2_. Site B/2 also resembles a bona-fide PI(4,5)P_2_ binding pocket (aromatic Y666 and Lys K684) (Fig. 7E).

Site C/4 has both Lys and aromatic residues but does not form a pocket and may be an electrostatic CALM-type PI(4,5)P_2_ binding site (Fig. 7F). Mutation of K480 abolishes the response to PI(4,5)P_2_, and mutation of R461 and R484 have a significant effect. However, mutation of the other three basic residues identified by MD has no effect.

In addition to these 3 major sites, mutagenesis identified several amino acids outside these sites. We suspect that some of these amino acids may play structural or allosteric roles. R579 makes a hydrogen bond with E568 in the loop between TM4 and TM5, which are important in forming the conduction pathway. R747 forms hydrogen bonds with L693 and E694 (L693:O-K747:NE and E694:O-K747:N), which lock the cytoplasmic ends of TM7 and TM8. This may be important in maintaining a proper conformation of Site B/2 (K682, R683, and R684) to interact with PI(4,5)P_2_. Mutation of R125 and its adjacent amino acids in the N-terminus significantly affect PI(4,5)P_2_ binding, but interpretation is hampered because the N-terminus is not modeled well in the cryo-EM structures (see below).

### Role of the N-terminus

It has been reported that ANO6, a paralog of ANO1 with 54% sequence similarity, is regulated by PI(4,5)P_2_ binding to the N-terminus (54, 55). Ye et al. (55) showed that mutation of amino acids K87, K88, K95, R96, K97, and R98 decreases the Ca^2+^-sensitivity and reduces the ability of exogenously-applied PI(4,5)P_2_ to restore ANO6 current after rundown. ANO6 residues 95-KRKR-98 align in PROMAL3D with ANO1 residues 124-KRFRR-128, and alignment of the ANO1 (5oyb) and ANO6 (6qp6) structures shows these sequences are similarly located in the protein structure. Although we find that mutation of these residues in ANO1 decreases the responsiveness to PI(4,5)P_2_, the N-terminus is not well resolved in the ANO1 structure, and amino acids 1-116 and 131-164 are not modeled. Thus, it remains unclear how these amino acids contribute to ANO1 regulation. Also, the role of these amino acids in ANO6 remains in question because Aoun et al. (54) showed that deletion or mutagenesis of this site in ANO6 had no effect on PI(4,5)P_2_ binding. Rather, they propose that amino acids 302-KKQPLDLIRK-311 in the proximal N-terminus of ANO6 are important for PI(4,5)P_2_ binding. This sequence aligns well with 313-KKQPLDLIRK-322 in ANO1, and we find that mutation of K313 in ANO1 significantly reduces the stimulatory effect of PI(4,5)P_2_ on ANO1 current. K313 is located in a re-entrant membrane loop just before TM1 in both ANO1 and ANO6 structures and is a crucial residue in Site A/1.

### A Network of PI(4,5)P_2_ Binding Sites

A major question raised by this study is “Why does ANO1 have multiple PI(4,5)P_2_ binding sites and what are their functions?” Most studies on PI(4,5)P_2_-protein interaction generally assume that the effect of PI(4,5)P_2_ is mediated by a single binding site. However, there are several examples of proteins interacting with PI(4,5)P_2_ in different ways. For example, X-ray crystallography, coarse-grained MD, and reconstitution in artificial membranes have demonstrated that the Arf GTPase and its GEF Brag2 form a complex that binds to the membrane via multiple PI(4,5)P_2_ interaction sites (65). There is also some mutational and biochemical evidence for multiple PI(4,5)P_2_ binding sites on profilin-1, gelsolin, TRPV1, EAG1, and K_ir_2.1 (66–70).

We anticipate that the three PI(4,5)P_2_ binding sites in ANO1 may have different and/or interacting effects. Although considerable work will be required to dissect these functional consequences, at this point in time we propose a general model for ANO1 gating that involves the binding of Ca^2+^ and PI(4,5)P_2_ to regulate this complex channel (Fig. 9). One corollary conclusion of this model is that the cryo-EM structures of ANO1 that have been published (22, 23) represent the inactivated state because the channel does not have PI(4,5)P_2_ bound.

**Figure 9.**
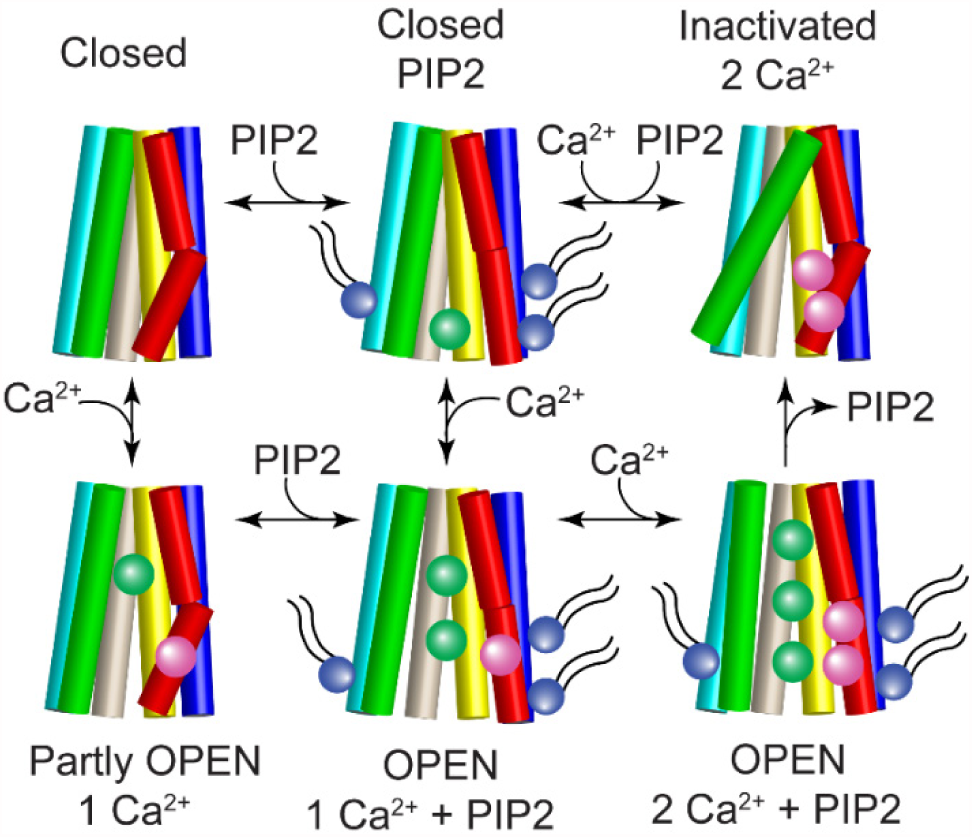
Cartoon model of ANO1 gating. TM2 (blue), TM3 (cyan), TM4 (green), TM5 (wheat), TM6 (red), and TM7 (yellow) are shown as cylinders. The pore is formed by TM4-TM7 and Ca^2+^ (magenta spheres) binds to residues in TM6 and TM7. When PI(4,5)P_2_ (purple sphere with tails) binds to the cytoplasmic ends of TM2 (Site A/1), TM6 (Site B/2), and TM3 (Site C/4), the cytoplasmic end of TM6 swings away from the pore to ultimately open the cytoplasmic vestibule to Cl^−^ (green spheres). Top row: the channel is closed in the absence of Ca^2+^ and inactivated when two Ca^2+^ ions are bound in the absence of PI(4,5)P_2_. Bottom row: the channel can partly open when one Ca^2+^ binds in the absence of PI(4,5)P_2_ but full channel opening requires both Ca^2+^ and PI(4,5)P_2_.

The locations of the three sites in the protein may provide some insights into their possible function. Key residues forming Site A/1 are located in TM1, TM2, and the loop between α0a and α0b. Because TM1 and TM2 are tightly packed against TM7 and TM8 that harbor part of the Ca^2+^ binding site, PI(4,5)P_2_ binding to Site A/1 could have widespread allosteric effects on Ca^2+^ sensitivity and could be responsible for stabilizing the Ca^2+^-bound open state. Site B/2 is formed by amino acids at the cytoplasmic end of TM6 and intracellular loop 3 **(ICL3)** connecting TM6 and TM7. TM6 plays a crucial role in ANO1 gating (24, 25, 30) so that its conformational changes could alter channel gating. Also, TM6 and TM7 harbor three amino acids that form the binding site for Ca^2+^, so PI(4,5)P_2_ binding to this site could also affect Ca^2+^-dependent gating. Site C/4 is located at the cytoplasmic end of TM3 in the loop connecting TM3 and TM2. TM3 is tightly packed with TM4 that lines the conduction pathway, so binding of PI(4,5)P_2_ to Site C/4 could modify the ion conduction pathway.

## Methods

### Cell Culture and Transfection

HEK-293 cells (ATCC) were maintained in modified DMEM with 10% FBS, 100 U/ml penicillin G and 100 *µ*g/ml streptomycin) and transiently transfected with mTMEM16A (Uniprot Q8BHY3). HEK293 cells were authenticated by short tandem repeat profiling and were tested for myoplasm contamination. TMEM16A was tagged on the C-terminus with EGFP. PCR-based mutagenesis was used to generate single amino acid mutations. Mutations were verified by sequencing.

### Phosphatidylinositols and Inositol Phosphates

Phosphatidylinositols were purchased from Echelon Research Laboratories (Salt Lake City, UT). Unless noted otherwise, the synthetic lipids used in these experiments have C8 saturated fatty acid chains. At concentrations used here, these short chain phosphoinositides are likely monodisperse in solution. It has been estimated that the critical micelle concentration (CMC) for lipids with phosphorylated inositol headgroups is >3 mM (71). Stock solutions of 10 mM PI(4,5)P_2_ were made in deionized H_2_O and stored at −20°C. Working solutions were made fresh immediately before the experiment.

### Preparation of the ANO1 Structure for Simulation

The cryo-EM structure of mouse TMEM16A (PDB 5OYB) at 3.75 Å resolution (72) was used as the starting structure for the MD simulations. Nomenclature of alpha-helices is from ref. (72). Because amino acids 1-116 (N-terminus), 131-164 (N-terminus), 260-266 (N-terminus), 467-487 (TM2-TM3 linker), 669-682 (TM6-TM7 linker), and 911-end (C-terminus) are unstructured in the cryo-EM model, we used a model in which the TM2-TM3 and TM6-TM7 linkers and N-terminal residues 131-164 and 260-266 were added using SuperLooper2 (73) and subjected to energy minimization. The two Ca^2+^ ions bound in each of the two subunits were preserved for all the simulations. The pKa of each ionizable residue was estimated using PROPKA (74, 75) and default protonation states were assigned based on the pKa analysis. Missing hydrogen atoms were added using PSFGEN in VMD (76). Internal water molecules were placed in energetically favorable positions within the protein using DOWSER (77, 78).

### Simulation System Setup

The ANO1 protein was first embedded in a palmitoyl-oleolyl phosphatidylcholine (POPC) bilayer generated using the CHARMM-GUI membrane builder (79). In each of the six independent simulation systems (Simu 1 – Simu 6), eight PI(4,5)P_2_ molecules were evenly placed around the protein in the inner leaflet of the membrane (∼1.4% PI(4,5)P_2_) (Fig. 5A). PI(4,5)P_2_ molecules were parameterized using CHARMM36 parameters. Considering the possible protonation states of PI(4,5)P_2_, two variations of PI(4,5)P_2_ headgroups were used in the simulations: systems 1–3 used palmitoyl-oleoyl-phosphatidylinositol-(4,5)-bisphosphate with protonation on P4, while systems 4–6 used the palmitoyl-oleoyl-phosphatidylinositol-(4,5)-bisphosphate with protonation on P5. The initial positions of the PI(4,5)P_2_ molecules were at least 30 Å away from any atom of the channel. The membrane was then converted to a highly mobile membrane memetic (HMMM) model using in-house scripts. This model replaces a portion of the membrane hydrophobic core by a more fluid representation using simple carbon solvent ethane (SCSE), while using short-tailed lipids to maintain full description of the headgroups and the initial part of the tails. The membrane/protein systems were fully solvated with TIP3P water (80) and buffered in 150 mM NaCl to keep the system neutral. The resulting systems consisting of ∼575,000 atoms were contained in a 244×180×141 Å^3^ simulation box.

To evaluate the specificity of PI(4,5)P_2_ binding to the channel, an additional set of simulations (3 trajectories, 350-ns each) were performed with 8 PtdSer molecules evenly placed around the protein in the inner leaflet of the membrane, in addition to the 8 PI(4,5)P_2_ molecules. Each simulation of mixed PI(4,5)P_2_ and PtdSer has a different initial placement of the lipids. The simulation setup was otherwise the same as described above.

### Simulation Protocols

MD simulations were carried out with NAMD2.12 (81) using CHARMM36m force field (82) and a timestep of 2 fs. Periodic boundary conditions (PBC) were used throughout the simulations. To evaluate long-range electrostatic interactions without truncation, the particle mesh Ewald (PME) method (83) was used. A smoothing function was employed for short-range non-bonded van der Waals forces starting at a distance of 10 Å with a cutoff of 12 Å. Bonded interactions and short-range non-bonded interactions were calculated every 2 fs. Pairs of atoms whose interactions were evaluated were searched and updated every 20 fs. A cutoff (13.5 Å) slightly longer than the non-bonded cutoff was applied to search for interacting atom pairs. Simulation systems were subjected to Langevin dynamics and the Nosé-Hoover Langevin piston method (84, 85) to maintain constant pressure (P=1 atm) and temperature (T=310 K) (NPT ensemble). To allow the simulation systems to fluctuate in all dimensions, in pressure control, a constant ratio was used for both HMMM and full-length simulations. This parameter keeps the x:y ratio of the unit cell constant rather than keeping the dimensions of the unit cell constant, thus allowing the surface area of the membrane to fluctuate during the simulation.

### HMMM Simulations

Because the HMMM representation simplifies the lipid bilayer by making the core more fluid, grid forces were applied on the carbon atoms affected by the conversion from the full-length membrane to the HMMM model to restrain the HMMM lipids, in order to resemble the full-length membrane while still allowing lipid molecules to fluctuate. The target atoms affected by the grid forces included carbon atoms in the solvent SCSE molecules and the terminal carbons (C26 and C36) of the short-tailed POPC and POPI lipids. The aim was to make the positions (z-coordinates) and flexibility of these target atoms resemble their counterparts in full-length membrane. To realize this, 3D grids with 1-Å spacing were specified in the simulation box to define the potential to be applied. Different potential values were then assigned to each grid point depending on the z-position within the membrane (centered at z=0). The grid potential *U*_*grid*_(*Z*) is defined by the equation,

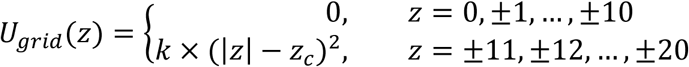

where 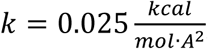, *z*_*c*_ = 10 Å, and *z* denotes the z-position of the grid point. For each target atom within the grid, the force was computed by a tri-cubic interpolation of the potential from the surrounding grid values based on the z-position of the atom. The target atoms thus experience zero or close-to-zero force when they are in the region comparable to that of their counterparts in the full-length membrane (z within ±10 Å); increasing force is experienced when they are diffusing toward the headgroup/bulk-solvent region (z beyond ±10 Å). Since only the terminal carbon atoms (C26 and C36) in the short-tail lipids are affected by the grid forces, this method provides the majority part of the molecules, especially the headgroups, with considerable flexibility in all dimensions during the simulation.

To verify that the HMMM simulations were providing a reasonable model of bilayer, we compared the dynamics of the phospholipids in a simulation of TMEM16A in full-length membrane (600 ns, no restraint) with the HMMM model used here (500 ns). The fluctuations of the choline-nitrogen atoms or phosphate atoms of POPC headgroups in the z-axis were used as metrics. In the HMMM simulations, the z-positions of the choline nitrogens within 10 Å of the protein fluctuate up to 10-10.5 Å in each direction, and phosphorus atoms can fluctuate 9-9.5 Å. The capacity for fluctuation of the headgroup components shows that the model used in this study provides sufficient freedom for the HMMM membrane mimetics to behave similarly to a full-length membrane (56, 64).

The six HMMM simulation systems were first energy-minimized for 10,000 steps followed by a 1-ns relaxation MD, during which the heavy atoms from the protein were positionally restrained (k=1 kcal/mol/Å^2^) to allow the *de novo* added structures to relax. Then a 500-ns equilibrium simulation was performed for each system with the Cα atoms of the transmembrane region (estimated using the PPM server) slightly restrained (k=0.1 kcal/mol/Å^2^). Our published data showed that using SCSE solvent can greatly improve the behavior of transmembrane proteins within the HMMM model (56). SCSE showed substantially less solvent intercalation into transmembrane proteins compared to DCLE and SCSM solvents in absence of restraint on the proteins (64). Nevertheless, to prevent any potential disruption of the transmembrane structure, we applied a slight restraint on the C-alpha atoms of the transmembrane region of TMEM16A. The rest of the protein (sidechains of the transmembrane region and the backbone atoms and sidechains of the cytoplasmic and extracellular regions) was free to move. Such restraint is not likely to affect the sampling of lipid-protein interactions because the transmembrane region of the channel does not move much during unrestrained simulations. We compared the dynamics of the protein in the HMMM simulations with the ones in normal membranes without restraint. The transmembrane region is stable and does not move much for either simulation (average heavy atom RMSD was 1.57 Å for HMMM vs 2.00 Å for full-length). The RMSD for the transmembrane region in full-length membranes over the measured time scale (600 ns) is only ∼0.43 Å larger than the HMMM simulations (500 ns). In comparison, the average heavy atom RMSD for the entire protein is 7.01 Å for HMMM simulations and 8.67 Å for full-length simulations, meaning that most of the flexibility of the protein originates from the dynamics of the cytoplasmic and extracellular domains. Based on this observation, we conclude that the slight restraint on the protein will not affect results of the simulations.

### Full-length Simulations

To investigate protein conformational changes upon PI(4,5)P_2_ binding, the PI(4,5)P_2_-bound HMMM systems were converted back to full-length phospholipids, followed by an additional 600-ns simulation without any restraints on the protein. A control simulation in the absence of PI(4,5)P_2_ molecules was also performed using the same protocol.

### Analysis of Lipid-protein Interactions

To characterize the interaction of lipid headgroups with potential binding sites on the protein, occupancy maps of the inositol group (and its phosphate groups) over the trajectories were calculated using the Volmap plugin in VMD (68). To determine the residues at those sites that stabilize PI(4,5)P_2_ binding, the polar interactions between the residues (side-chain nitrogen/oxygen atoms) and the PI(4,5)P_2_ inositol ring (phosphate/hydroxyl groups) in the binding site were calculated for each frame of the trajectory using a distance cutoff of 4 Å for phosphorus atoms of the phosphates and 3.5 Å for oxygen atoms of the hydroxyls.

### Statistical Analysis

Data were analyzed using Origin 2017 SR2. Error bars are standard error of the mean. Statistics are described in each figure.

### Data Sharing

All data presented in this paper will be made available to investigators upon reasonable request.

## Acknowledgements

Research reported in this publication was supported by the National Eye Institute, National Institute of Arthritis and Musculoskeletal Diseases, and National Institute of General Medical Sciences of the National Institutes of Health under award numbers R01EY114852 (to HCH), R01AR067786 (to HCH), P41-GM104601 (to ET), and R01-GM123455 (to ET). 95% of this research was financed with Federal money and 5% from non-governmental sources. The content is solely the responsibility of the authors and does not necessarily represent the official views of the National Institutes of Health. We also acknowledge computing resources provided by Blue Waters at National Center for Supercomputing Applications, and Extreme Science and Engineering Discovery Environment (grant MCA06N060 to ET).

